# Temporal genomic erosion estimates require demographic and technical validation

**DOI:** 10.64898/2026.11.24.768837

**Authors:** Michael V. Westbury, Hernán E Morales

**Author notes:** Corresponding author: Michael V Westbury.

## Abstract

Genetic diversity is a fundamental component of biodiversity, and temporal genomics offers a powerful approach to monitoring genomic erosion and informing conservation assessments. However, robust inference of genetic diversity loss requires consistency with demographic expectations and careful evaluation of technical biases between historical and modern sequencing data. Here, we combine population-genetic simulations, independent reanalyses of temporal genomic datasets, comparisons of alternative heterozygosity estimators and controlled experiments of historical-like DNA degradation to evaluate the reliability of genomic erosion estimates. Using data from four bird species included in a recent study reporting average diversity losses of 43–49% across 62 species (Liu et al., 2026), we show that losses of this magnitude generally require severe, sustained population bottlenecks accompanied by extensive runs of homozygosity (ROH), which were uncommon in the modern genomes examined. Heterozygosity estimates were highly sensitive to analytical methods: ANGSD recovered substantial declines (23–42%), whereas HEARTY and ATLAS showed little or no loss (−1% to +9%). Experimentally introducing historical-like DNA degradation into reads from genetically identical individuals generated apparent declines of up to 42% with ANGSD, demonstrating that sequencing artefacts alone can produce signals comparable to those reported. We further evaluated estimator performance using the Chatham Island black robin, which experienced a documented severe demographic collapse. HEARTY and ATLAS recovered substantial heterozygosity losses (56–59%), although additional degradation experiments revealed dataset-specific biases in both methods. Together, our findings demonstrate that technical artefacts can mimic or obscure genuine genomic erosion, with effects varying across datasets and analytical approaches. Integrating demographic expectations, complementary genomic indicators and explicit assessments of sequencing biases is essential for reliable use of temporal genomics in biodiversity monitoring and conservation assessments.

## Introduction

Genetic diversity is an essential component of biodiversity, and temporal genomics provides a powerful approach to assessing genomic erosion and informing conservation decisions ^1–3^. However, the magnitude of diversity loss depends on demographic history, effective population size, population connectivity and the time elapsed since demographic decline, which are not always known. Furthermore, systematic differences in DNA preservation, sequencing quality and analytical methodology between ancient/historical and modern samples can introduce biases that mimic or obscure genuine genomic change ^4,5^. Reliable interpretation of temporal genomic data therefore requires evaluating both the demographic plausibility of inferred changes and their sensitivity to technical artefacts.

A recent study by Liu et al. (2026) ^6^ assembled an extensive temporal genomic dataset spanning 62 bird species, reporting average losses of 43–49% of genomic diversity over a median interval of 49 years (approximately 20 generations). Such substantial losses over relatively few generations are extraordinary, particularly in species without documented severe demographic collapses. These findings raise important questions about whether the inferred changes are consistent with plausible demographic histories and whether they are robust to differences in historical and modern sequencing data.

Here, we investigate these questions using complementary approaches. First, we use forward-time simulations to evaluate the demographic conditions required to produce substantial heterozygosity loss and the accompanying genomic signatures. Next, we reanalyse historical and modern genomes from four species investigated by Liu et al., comparing alternative heterozygosity estimators and testing whether simulated historical-like DNA degradation can generate apparent diversity declines. Finally, we reanalyse temporal genomic data from the Chatham Island black robin (*Petroica traversi*)^7^, which experienced a well-documented severe population bottleneck. Together, these analyses examine how demographic expectations and dataset-specific technical biases influence the interpretation of genomic erosion, highlighting the importance of independently validating temporal genomic estimates.

## Results and discussion

### Severe demographic bottlenecks are required to observe large losses of genetic diversity

To assess the demographic conditions required to reproduce the magnitude of genomic erosion reported by Liu et al. (2026), we used SLiM to simulate five interconnected populations undergoing demographic decline and reduced migration over 10–20 generations. Populations initially comprised 1,000 individuals per deme before declining to 10–100 individuals, with migration rates ranging from 0 to 10%. Heterozygosity loss increased with bottleneck severity and duration, consistent with neutral population genetic expectations^8^ (Fig. 1A). For example, under isolation, an effective population size of 20 is expected to produce approximately 40% heterozygosity loss over 20 generations, compared with less than 10% for an effective population size of 100.

**Figure 1.**
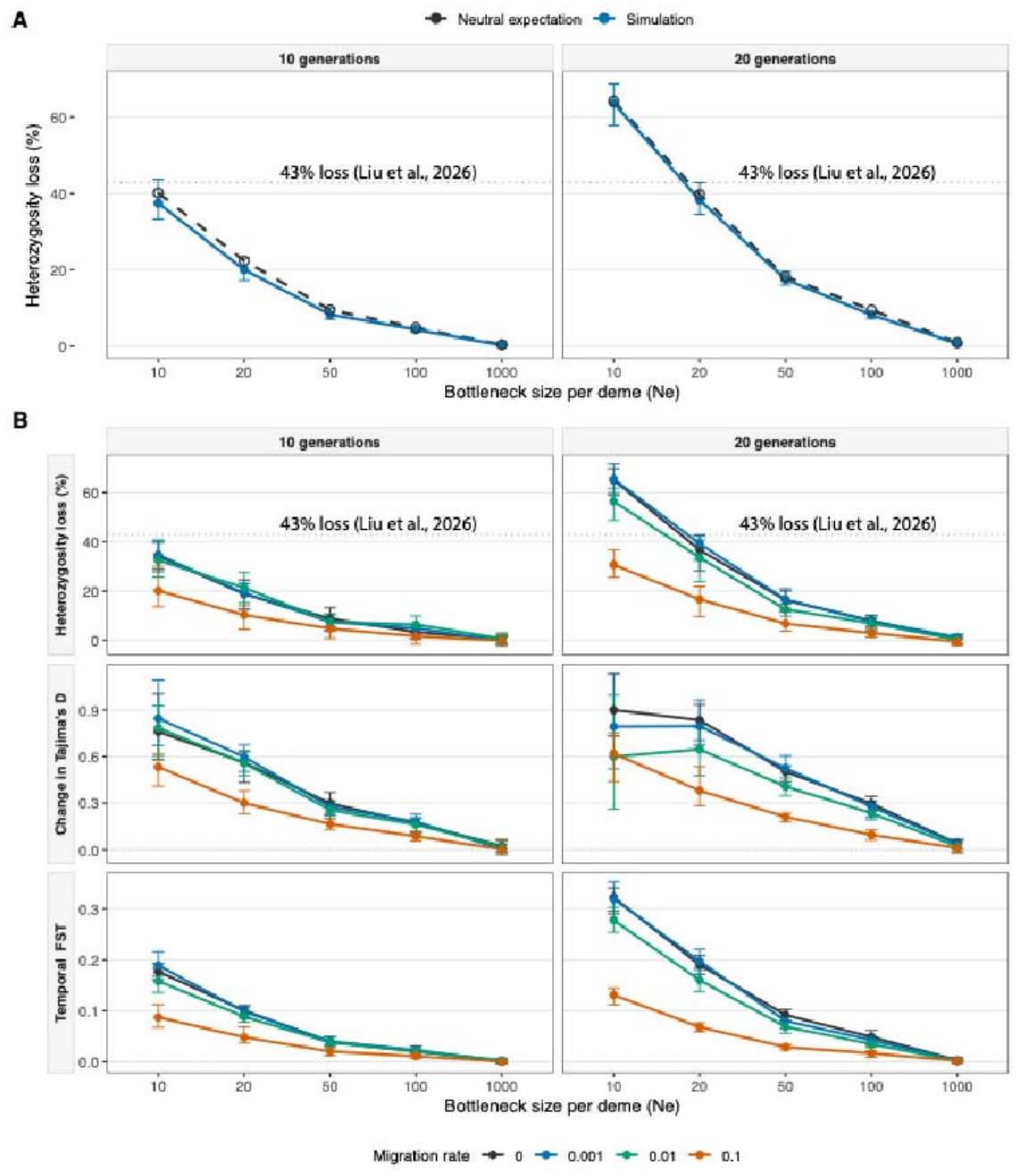
Theoretical and simulated expectations of demographic decline on genomic diversity over 10 and 20 generations. Forward-time simulations of five interconnected demes undergoing changes in population size and migration. **(A)** Comparison of theoretical neutral expectations (black dashed lines) and simulated heterozygosity loss (blue lines) under complete isolation. The expected proportion of heterozygosity retained after *t* generations was calculated as (H_t/H_0=(1-1/(2Ne))^t) for bottlenecks of 10 and 20 generations. **(B)** Effects of post-decline population size (*Ne* = 10, 20, 50, 100 or 1,000) and migration rate (*m* = 0, 0.001, 0.01 or 0.1) on heterozygosity loss, change in Tajima’s D, and temporal genetic differentiation (Hudson’s FST) between historical and modern populations. Demes initially comprised 1,000 individuals, with *N* = 1,000 representing the no-decline scenario.

Simulated losses approaching the 43–49% reported by Liu et al. were reproduced primarily under severe population bottlenecks, whereas more moderate reductions produced substantially smaller losses, even when connectivity was restricted (Fig. 1B). Our simulations also reproduced pronounced changes in Tajima’s D and temporal FST under severe demographic declines, with these responses generally attenuated by higher migration rates (Fig. 1B). These results indicate that fragmentation alone does not necessarily generate rapid, substantial genomic erosion^9^. Importantly, census and effective population sizes are not equivalent, and their relationship varies among species and demographic conditions^10,11^. Nevertheless, the simulations provide a quantitative benchmark for assessing whether inferred genomic changes are consistent with plausible demographic histories. Although Liu et al. attribute the reported erosion primarily to fragmentation, demographic collapses of the severity required by our simulations have not been independently established for their study species.

Severe bottlenecks in our simulations also produced increased genomic homozygosity, including longer homozygous tracts (Fig. 2A). We therefore examined runs of homozygosity (ROH) in modern genomes from four species investigated by Liu et al.: *Fringilla montifringilla, Actinodura cyanouroptera, Trochalopteron erythrocephalum* and *Yuhina occipitalis*. Long ROH (>1 Mb) were uncommon, and no individual exhibited segments exceeding 5 Mb (Fig.2B). Although these findings do not exclude historical bottlenecks or more gradual reductions in effective population size, they provide limited evidence of extensive recent inbreeding. Heterozygosity and ROH reflect demographic processes over different timescales, emphasising the value of interpreting genomic erosion alongside complementary demographic indicators.

**Figure 2.**
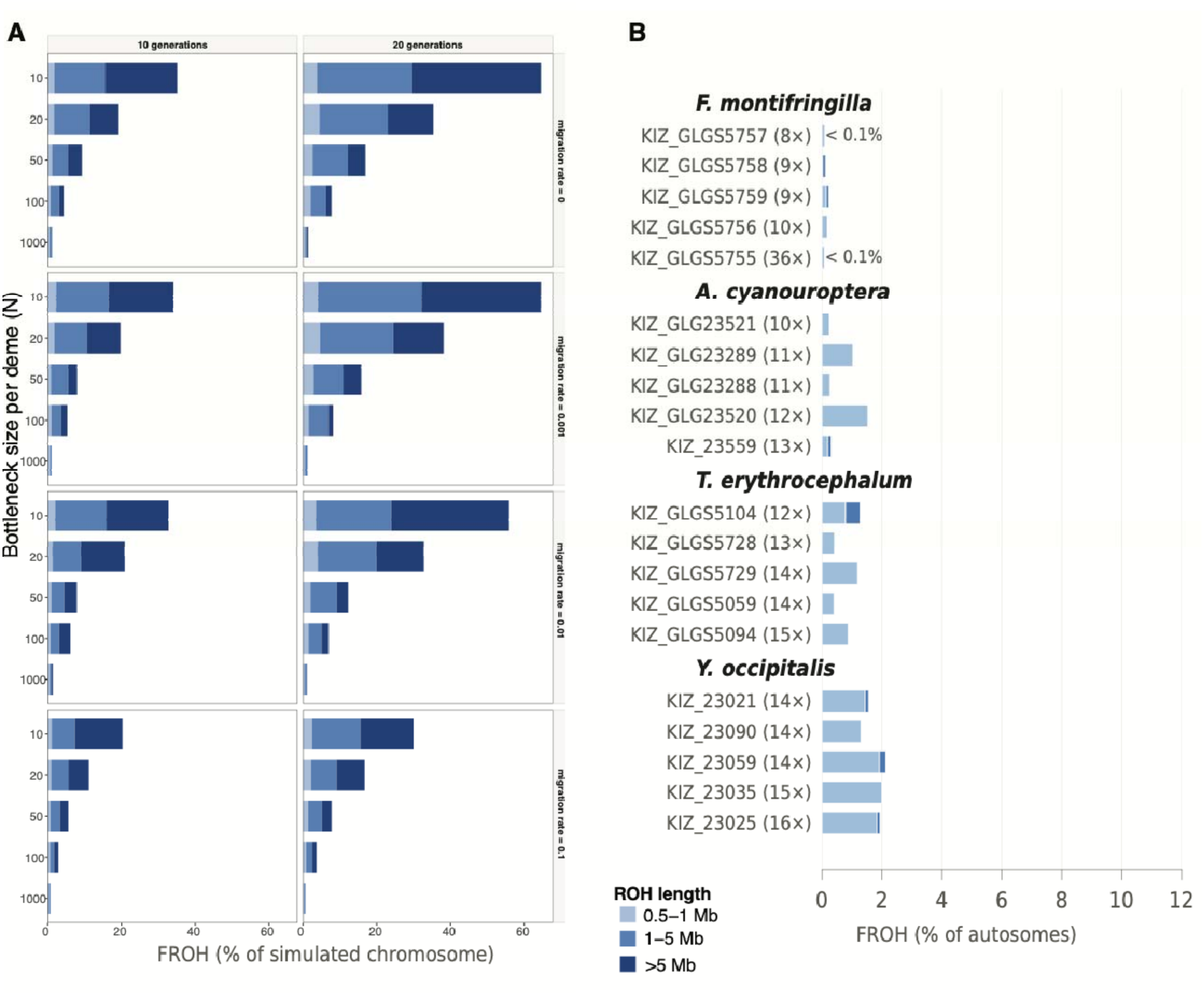
Simulated and empirical runs of homozygosity (ROH). **(A)** Simulated genomic homozygosity (FROH) under varying bottleneck sizes, migration rates and durations. **(B)** FROH in modern individuals from four species investigated by Liu et al. (2026), estimated using BCFtools. Stacked bars indicate the fraction of the genome within homozygous tracts of 0.5–1 Mb, 1–5 Mb and >5 Mb, with darker shading representing longer tracts.

### Temporal heterozygosity estimates are highly sensitive to analytical method

We independently reanalysed five historical and five modern genomes from each of the four species (Fig. 3A). Although Liu et al. reported heterozygosity declines of 30–57%, our ANGSD analyses recovered declines of 26–42% at native sequencing depth. Matching historical and modern coverage and applying mapDamage base-quality rescaling reduced these estimates to 23–37%. Restricting analyses to transversions likewise failed to consistently eliminate the apparent declines. In contrast, HEARTY, which standardises heterozygous-site detection through read sampling and alternative-allele support, and ATLAS, which explicitly models sequencing errors and post-mortem damage, recovered little evidence of heterozygosity loss across the four species (−1% to +9%; Fig. 3). These discrepancies raise concerns about the robustness of the original estimates.

**Figure 3.**
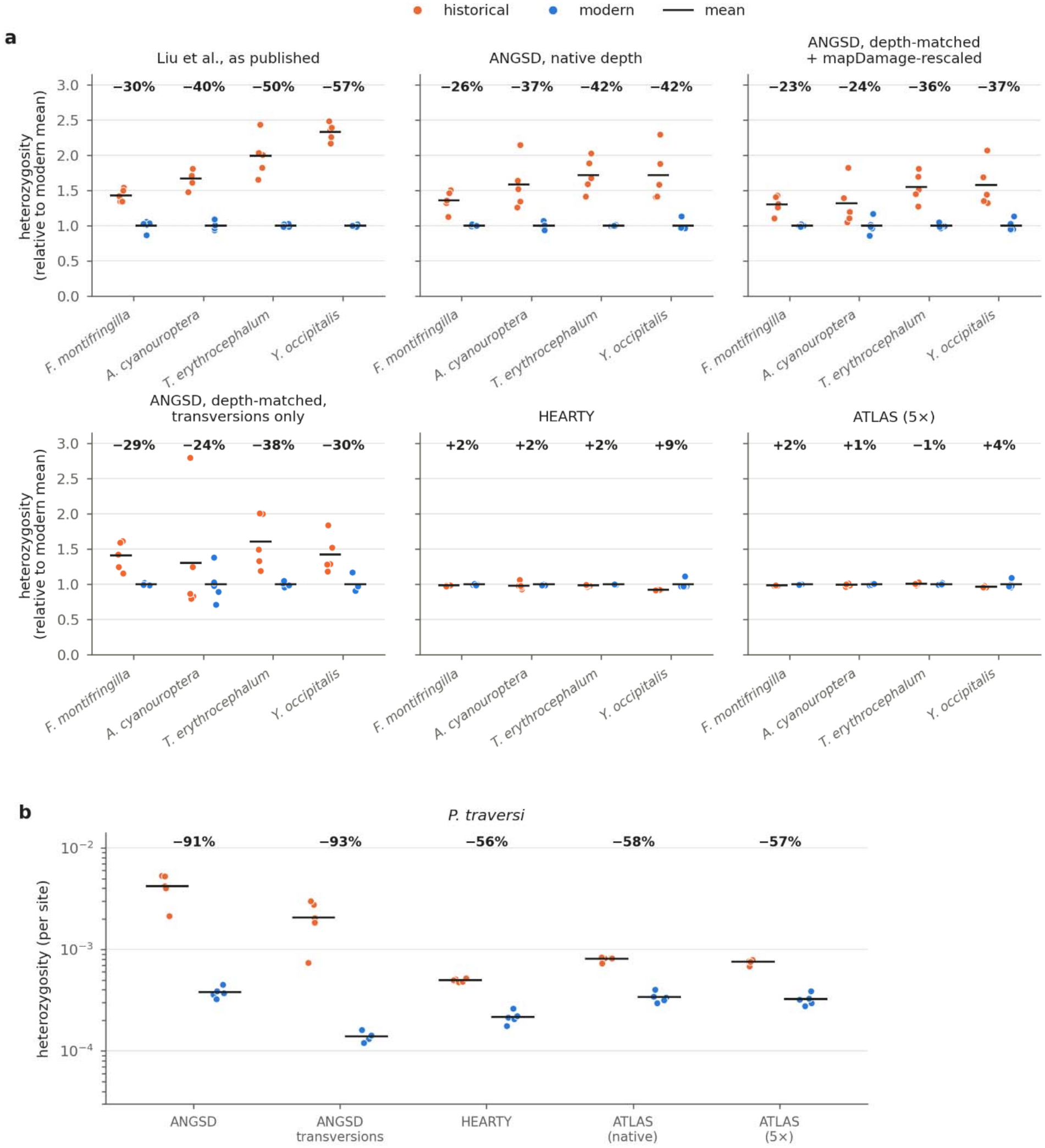
Sensitivity of temporal heterozygosity estimates to analytical method across bird species. **(A)** Historical and modern heterozygosity estimates for four bird species from Liu et al. (2026), comparing published estimates with ANGSD (native depth, depth-matched and damage-rescaled, and transversions only), HEARTY and ATLAS. Values are normalised to the corresponding modern mean for each species and method. **(B)** Heterozygosity estimates for the Chatham Island black robin (*Petroica traversi*), shown per site on a logarithmic scale. Orange and blue points represent historical and modern individuals, respectively; horizontal black lines indicate period means. Percentages indicate temporal changes in mean heterozygosity, with negative values representing declines.

To investigate the source of these discrepancies, we examined substitution patterns in the sequencing reads. Relative to doubleton mismatches, historical samples exhibited greater enrichment of singleton C→T/G→A transitions (2.5–8.9-fold) and G→T/C→A transversions (1.1–7.5-fold) than modern samples (Fig. S1). However, four of five modern *A. cyanouroptera* individuals also exhibited a pronounced G→T/C→A excess, demonstrating that such mismatch patterns are not restricted to historical material. Similar transversion excesses have previously been documented in historical specimens^12^. These patterns suggest that erroneous single-read observations may contribute to inflated heterozygosity estimates and that excluding transitions alone may be insufficient to eliminate technical biases.

To determine whether the observed mismatch patterns could explain the apparent heterozygosity declines, we used SCAR to introduce historical-like fragmentation, deamination and exces transversion mismatches into sequencing reads from three modern *F. montifringilla* individuals. Treating degraded reads as historical and their unmodified counterparts as modern, ANGSD recovered an apparent 25% heterozygosity decline, approaching the 30% reported by Liu et al. for this species (Fig. 4A). Restricting the analysis to transversions increased the apparent decline to 42%, whereas HEARTY and ATLAS estimates remained largely unchanged (+3% and +2%, respectively). Because the underlying genomes were identical, these differences arose entirely from technical artefacts, demonstrating that historical-like sequencing errors can generate apparent genomic erosion comparable to the original estimates.

**Figure 4.**
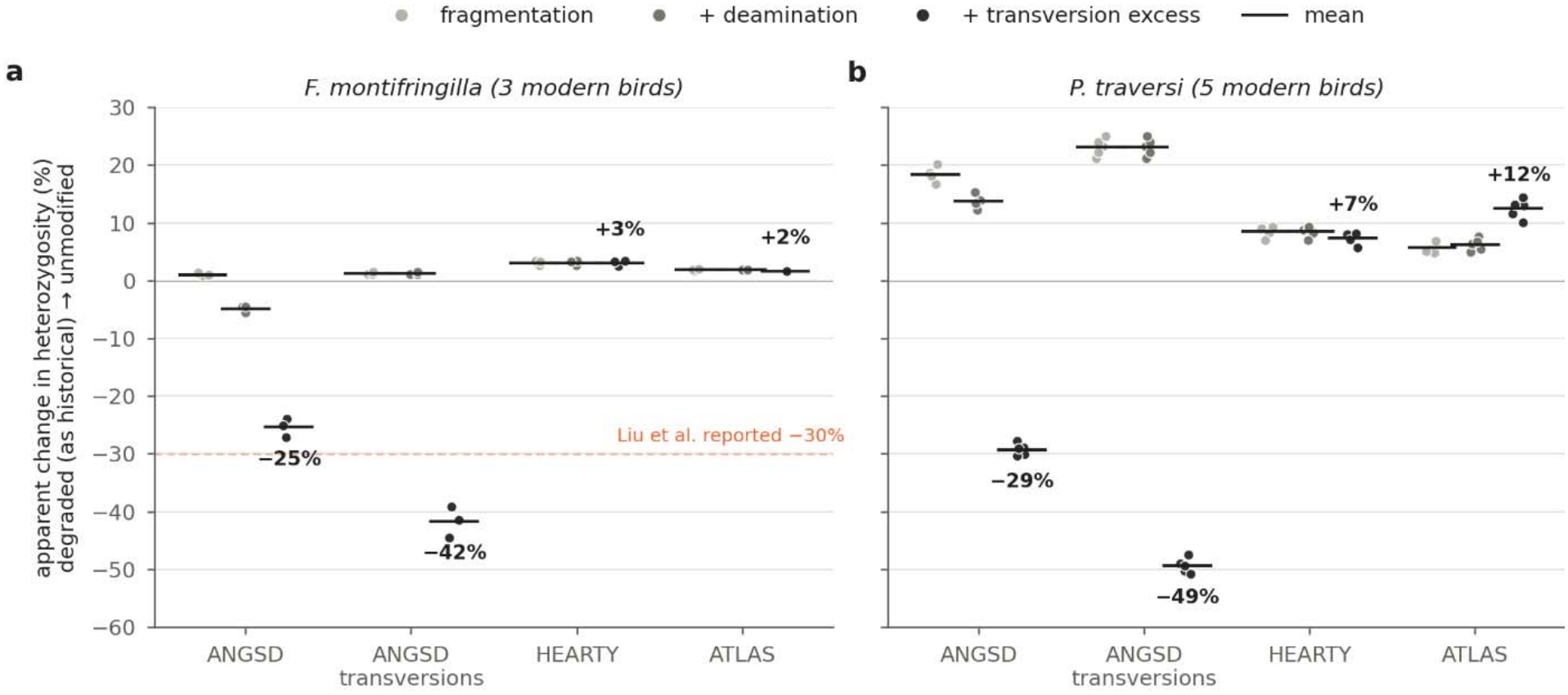
Effects on heterozygosity estimates of experimentally introduced historical-like sequencing degradation into modern reads. Modern sequencing reads from **(A)** *Fringilla montifringilla* (n = 3) and **(B)** *Petroica traversi* (n = 5) were progressively degraded using SCAR through fragmentation, terminal deamination and excess G→T/C→A mismatches. Points represent individual datasets following each cumulative treatment, and horizontal black line indicate means. Percentage changes were calculated as 100 × (H_unmodified − H_degraded)/H_degraded, treating degraded reads as pseudo-historical. The dashed orange line in (A) indicates the −30% decline reported by Liu et al. (2026).

To investigate whether alternative tools could also recover substantial genomic erosion, we reanalysed temporal genomes from the Chatham Island black robin (*Petroica traversi*), which experienced a documented population collapse to five individuals, including a single successfully breeding pair. Previous genomic analyses revealed substantial heterozygosity loss and extensive long ROH following this bottleneck^7^, providing independent evidence of severe demographic collapse.

Using five historical and five modern genomes, HEARTY and ATLAS recovered heterozygosity declines of approximately 56% and 57–59%, respectively, consistent with the robin’s documented demographic history (Fig. 3B). In contrast, conventional ANGSD suggested an inflated 91% decline. Similar to the Liu et al dataset, historical reads exhibited elevated mismatches across multiple substitution classes (Fig. S1).

However, controlled degradation experiments demonstrated that HEARTY and ATLAS were not entirely insensitive to technical differences in this dataset (Fig. 4B). Treating degraded reads as pseudo-historical and unmodified reads as modern, HEARTY and ATLAS recovered apparent heterozygosity increases of approximately 7% and 12%, respectively, whereas ANGSD recovered an apparent decline of 29%. Thus, degradation could mask genuine genomic erosion under HEARTY and ATLAS while exaggerating it under ANGSD. These effects differed substantially from those observed in *F. montifringilla*, highlighting how specimen preservation, sequencing characteristics and analytical assumptions can influence both the magnitude and direction of estimation bias. Although the alternative estimators recovered substantial genomic erosion consistent with the robin’s demographic history, neither can be assumed to provide universally unbiased estimates.

### Implications for temporal genomic inference

Together, these findings illustrate the value of combining demographic and sequencing simulations when assessing temporal genomic change. Forward-time simulations, such as those implemented in SliM^13^, evaluate whether inferred diversity losses are consistent with plausible demographic histories, while read-level degradation experiments, such as those implemented in SCAR, establish whether technical differences can independently generate or obscure comparable signals. Importantly, the contrasting results from the four species reanalysed from Liu et al. and the black robin demonstrate that the magnitude and direction of estimation biases can differ substantially among datasets. Methods that recover substantial genomic erosion consistent with a documented demographic collapse can nevertheless remain somewhat sensitive to technical artefacts. Consequently, neither agreement among estimators nor the application of standard damage corrections necessarily establishes that an estimate is unbiased. Both approaches are conditional on their underlying assumptions but provide complementary assessments of uncertainty that cannot necessarily be resolved through quality filtering or alternative heterozygosity estimators alone.

The dataset assembled by Liu et al. represents an important resource for conservation genomics, but the discrepancies identified in our four-species reanalysis warrant a broader reassessment of its reported diversity losses. More generally, genetic risk extends beyond changes in genome-wide heterozygosity to include inbreeding, loss of adaptive variation, immunogenetic diversity and accumulation of deleterious alleles ^14–17^. We therefore support incorporating genomic indicators into biodiversity monitoring, while advocating for routine evaluation of demographic plausibility, dataset-specific sequencing biases, complementary genomic evidence, and harmonised methods and data-reporting practices. Reliable assessments of genomic erosion require distinguishing genuine biological change from artefacts of its measurement.

## Materials and Methods

### Study design and source data

The four-species empirical analyses used five historical and five modern individuals per species, except where estimator-specific coverage criteria required exclusion. For full computational reproducibility, exact model versions, random seeds and analysis scripts will accompany the preprint.

### Forward-time demographic simulations and theory

We conducted forward-time simulations in SLiM 4 ^13^ using a neutral, diploid Wright–Fisher model with non-overlapping generations and five interconnected demes. The simulated genomic architecture was based on collared flycatcher chromosome 12, retaining annotated exonic, intronic and intergenic regions and an empirical recombination map^18^. Neutral mutations occurred at a rate of 5 × 10□□ per site per generation. Following a 25,000-generation burn-in with Ne = 1,000 individuals per deme and symmetric island migration (m = 0.1), we simulated abrupt changes in population size and migration lasting 10 or 20 generations. The simulation grid included per-deme population sizes of Ne = 10, 20, 50, 100 and 1,000 and migration rates of 0, 0.001, 0.01 and 0.1, representing the total proportion of offspring derived from parents in other demes.

We performed 20 independent replicates per scenario, sampling five individuals per deme before and after demographic change. We calculated heterozygosity, changes in Tajima’s D, temporal Hudson’s FST and individual runs of homozygosity (ROH). ROH were identified as homozygous intervals ≥0.5 Mb based on simulated heterozygous markers and summarised by length class and FROH. The no-migration expectation for retained neutral heterozygosity was calculated using the standard diploid drift expression H□/H□ = (1−1/(2N□))□. We compared simulation means and variability with this expectation for each bottleneck size and duration.

### Sequencing data and read processing

We selected four of the 62 species investigated by Liu et al. (2026): *Fringilla montifringilla, Actinodura cyanouroptera, Trochalopteron erythrocephalum* and *Yuhina occipitalis*.. *F. montifringilla* is a winter visitor to the region that breeds across the Palearctic, making a ~30% loss of heterozygosity within ~20 generations through local habitat change implausible; it therefore serves as a natural negative control. It also had the highest historical heterozygosity of the 62 species and has a chromosome-level reference assembly available. *A. cyanouroptera* had the shortest historical insert sizes in the dataset (median 61 bp), representing the most fragmented samples. *Y. occipitalis* had among the lowest heterozygosity and one of the largest reported declines (57%), and *T. erythrocephalum* showed a similarly large reported decline (50%). Together, the four species span the range of baseline diversity, DNA fragmentation and reported heterozygosity loss in the original study.

We downloaded raw paired-end reads from the Genome Sequence Archive (historical: CRA032748; modern: CRA020079). We mapped *F. montifringilla* to the chromosome-level bFriMon1 assembly (GCA_988331165), and *T. erythrocephalum* and *Y. occipitalis* to their species-specific assemblies from Liu et al. (2026) (KIZ_MT275 and KIZ_GLGS1218), Reads from our *A. cyanouroptera* samples differed from the published *A. cyanouroptera* assembly (KIZ_GLGS6485) at ~9% of aligned bases, compared with ~2.5% for *A. egertoni*, raising concerns about the taxonomic assignment of the former. We therefore used the *A. egertoni* assembly as the reference for these samples (KIZ_GLGS5798).

We processed historical and modern sequences using PlainMap v0.1.0 ^19^. We trimmed adapters and merged overlapping read pairs with fastp v1.0.1 ^20^, retaining reads of at least 30 bp. We aligned reads using BWA-MEM v0.7.17 ^21^, retained alignments with mapping quality ≥20 and removed PCR duplicates using samtools markdup. We restricted subsequent analyses to autosomes, excluding sex chromosomes and unplaced scaffolds. Mean autosomal sequencing depth ranged from 4.8–12.0× in historical individuals and 7.9–36.3× in modern individuals.

We additionally rescaled base qualities using mapDamage v2.2.3^22^. We estimated damage profiles independently for each individual using two million randomly selected reads and applied the resulting rescaling models to the complete alignments.

### Individual heterozygosity estimation

We estimated individual heterozygosity using ANGSD, HEARTY and ATLAS to evaluate the influence of sequencing depth, post-mortem damage and analytical methodology.

#### ANGSD

We estimated heterozygosity using ANGSD v0.935 ^22^ with the GATK genotype-likelihood model (-GL 2), minimum mapping and base qualities of 30, uniquely mapped reads and removal of problematic alignments. We retained sites with depths between 0.5 and 2 times each individual’s mean autosomal coverage. We estimated heterozygosity from the heterozygous class of the individual folded site frequency spectrum using realSFS. We performed analyses using all substitutions and transversions only (-noTrans 1).

To evaluate coverage-related bias, we ranked historical and modern individuals by sequencing depth within each species and paired individuals accordingly. We downsampled modern alignments to match their historical counterparts using samtools view --subsample and recalculated the depth filters. We repeated these analyses using mapDamage-rescaled alignments for both sampling periods.

#### HEARTY

We estimated individual heterozygosity using HEARTY (commit d8305fe; https://github.com/BiodiversityExtinction/HEARTY), which identifies heterozygous sites from nucleotide counts using a user-defined minor-allele frequency threshold. We generated per-site A, C, G and T counts using ANGSD v0.935 (-doCounts 1 -dumpCounts 4 -uniqueOnly 1 - remove_bads 1), requiring minimum base and mapping qualities of 30 (-minq 30 -minmapq 30). We retained autosomal sites with coverage between five reads and twice each individual’s mean sequencing depth.

To standardise heterozygosity estimation across individuals with different sequencing depths, we used HEARTY’s exact downsampling mode (--downsample-depth 5 --downsample-mode exact) with a minor-allele frequency threshold of 0.25 (-t 0.25). At a standardised depth of five reads, this threshold requires at least two reads supporting the minor allele for a heterozygous classification. For sites with more than five reads, HEARTY calculates the probability of a heterozygous call across all possible five-read combinations, weighted by their multivariate hypergeometric probabilities. Each site therefore contributes its expected heterozygous-call probability rather than a binary classification based on a randomly selected subset of reads. We calculated genome-wide heterozygosity as the sum of these expected probabilities divided by the number of eligible sites.

We performed analyses using all substitutions and transversions only, with both original and mapDamage-rescaled alignments.

#### ATLAS

We estimated heterozygosity using ATLAS v2.0.4 ^23^, which explicitly models sequencing errors and post-mortem damage. We selected calibration sites using GERP conservation scores from the Ensembl release 115 alignment of 27 sauropsid genomes ^24^. We used canary (*Serinus canaria*) as the source genome for *F. montifringilla* and great tit (*Parus major*) for the three babbler species. We lifted the 20 million highest-scoring sites to each reference genome and retained the 10 million highest-scoring autosomal sites for calibration.

For each individual, we used estimateErrors to fit sample-specific damage and base-quality recalibration models incorporating read position, sequence context, fragment length and mapping quality. We treated the calibration sites as invariant during model fitting. We then estimated heterozygosity under the HKY85 substitution model using summaryStats, sampling 1 Mb windows at 10 Mb intervals across the autosomes. We excluded calibration sites and applied minimum mapping quality of 30, base qualities of 20–93, a maximum of 10% soft-clipped bases and a maximum of 5% mismatches per read.

We standardised samples to 5× coverage for historical–modern comparisons. We excluded two historical individuals with insufficient coverage (*A. cyanouroptera* KIZ_003002 and *Y. occipitalis* KIZ_010911).

### Runs of homozygosity

We inferred ROH in the five modern individuals from each of the four species using bcftools roh ^25^. We generated autosomal genotypes using minimum mapping and base qualities of 30 and retained high-quality biallelic variants with site quality ≥30 and depths between 0.5 and 2 times each individual’s mean coverage. We estimated allele frequencies separately for each species using its five modern individuals.

We classified ROH into three length categories (0.5–1 Mb, 1–5 Mb and >5 Mb) and calculated the fraction of the autosomal genome contained within ROH (FROH). We restricted these analyses to modern individuals because the historical sequences were more susceptible to sequencing-error-induced false heterozygous calls.

### Characterisation of substitution mismatch spectra

To investigate potential sources of heterozygosity estimation bias, we characterised single-read and double-read non-reference observations using HEARTY nucleotide counts. We sampled five reads per site using a fixed random seed and classified sites containing exactly one or two reads supporting the same non-reference allele by substitution type.

We compared C→T and G→A transitions with their corresponding reverse substitutions (T→C and A→G), and G→T and C→A transversions with G→C and C→G controls, respectively. We also generated position-specific damage profiles using mapDamage and quantified the excess of G→T/C→A mismatches relative to their corresponding control substitutions.

### Experimental introduction of historical-like degradation

We used SCAR ^4^ to investigate whether historical-like differences in fragment length and mismatch profiles could independently generate apparent changes in heterozygosity. We selected three modern *F. montifringilla* individuals (KIZ_GLGS5756, KIZ_GLGS5757 and KIZ_GLGS5758) and progressively introduced degradation into their sequencing reads.

After adapter trimming and read merging, we reverse-complemented unmerged reverse reads to produce consistently oriented single-end sequences. We introduced degradation cumulatively using three empirically derived historical profiles: (i) fragmentation matching the read-length distribution of KIZ_013086 (mean 145 bp); (ii) position-specific terminal C→T and G→A substitutions based on KIZ_013087, which exhibited the strongest deamination signal; and (iii) G→T and C→A substitutions based on KIZ_013083, which exhibited the greatest excess of these mismatches relative to corresponding control transversions.

We independently mapped each treatment using the same pipeline as the empirical data and compared the resulting fragment-length and mismatch profiles with their historical targets. We downsampled treatments to approximately 4.7× coverage for ANGSD comparisons and estimated heterozygosity using ANGSD, HEARTY and ATLAS as above.

Because the degraded and unmodified reads originated from the same individuals, their underlying genetic diversity was identical. To express technical differences as apparent historical–modern genomic change, we treated degraded reads as historical and unmodified reads as modern and calculated percentage change as 100 × (H_unmodified − H_degraded)/H_degraded.

### Chatham Island black robin reanalysis

We additionally reanalysed whole-genome sequencing data from five historical and five modern Chatham Island black robins (ENA PRJEB46470)^7^. Historical sequences were available as previously trimmed and merged single-end reads, whereas modern sequences were provided as trimmed paired-end reads. We mapped sequences to the *P. traversi* reference genome (GCA_025920805.1) using PlainMap and restricted analyses to the 99 autosomal scaffolds.

We applied the same ANGSD, HEARTY and ATLAS heterozygosity estimation procedures described above, using great tit as the source genome for ATLAS calibration. We analysed the original alignments and additionally assessed the effects of depth matching, mapDamage rescaling and transversion-only filtering. We evaluated ATLAS estimates at several standardised sequencing depths, including 5×.

To investigate the effects of historical-like degradation, we applied SCAR to reads from all five modern black robins using the fragment-length and mismatch profiles of historical individual BRH08 (mean fragment length 71 bp). We introduced fragmentation, terminal deamination and excess G→T/C→A mismatches cumulatively, following the procedure used for *F. montifringilla*. We analysed the resulting datasets using ANGSD, HEARTY and ATLAS, limiting ANGSD comparisons to approximately 6× and ATLAS to no more than 5× coverage while retaining full eligible depth for HEARTY.

For these experiments, we treated degraded reads as pseudo-historical and their unmodified counterparts as modern, calculating apparent temporal changes as 100 × (H_unmodified − H_degraded)/H_degraded, consistent with the *F. montifringilla* experiments and empirical temporal comparisons.

### Temporal comparisons

We calculated empirical temporal changes in mean heterozygosity following Liu et al. (2026), using: %ΔH = 100 × (H_modern − H_historical)/H_historical. Negative values indicate heterozygosity loss, whereas positive values indicate increased heterozygosity. We summarised changes using individual heterozygosity estimates within each sampling period.

## Data and code availability

The sequencing datasets analysed in this study are publicly available through the Genome Sequence Archive (GSA; accessions CRA020079 and CRA032748) and the European Nucleotide Archive (ENA; accession PRJEB46470). Analysis scripts, SLiM simulation code, SCAR degradation profiles, sample metadata, processed summary data and underlying figure data are available at https://github.com/Mvwestbury/TemporalGenomicsEvaluation.

## Supplementary Information

**Figure S1.**
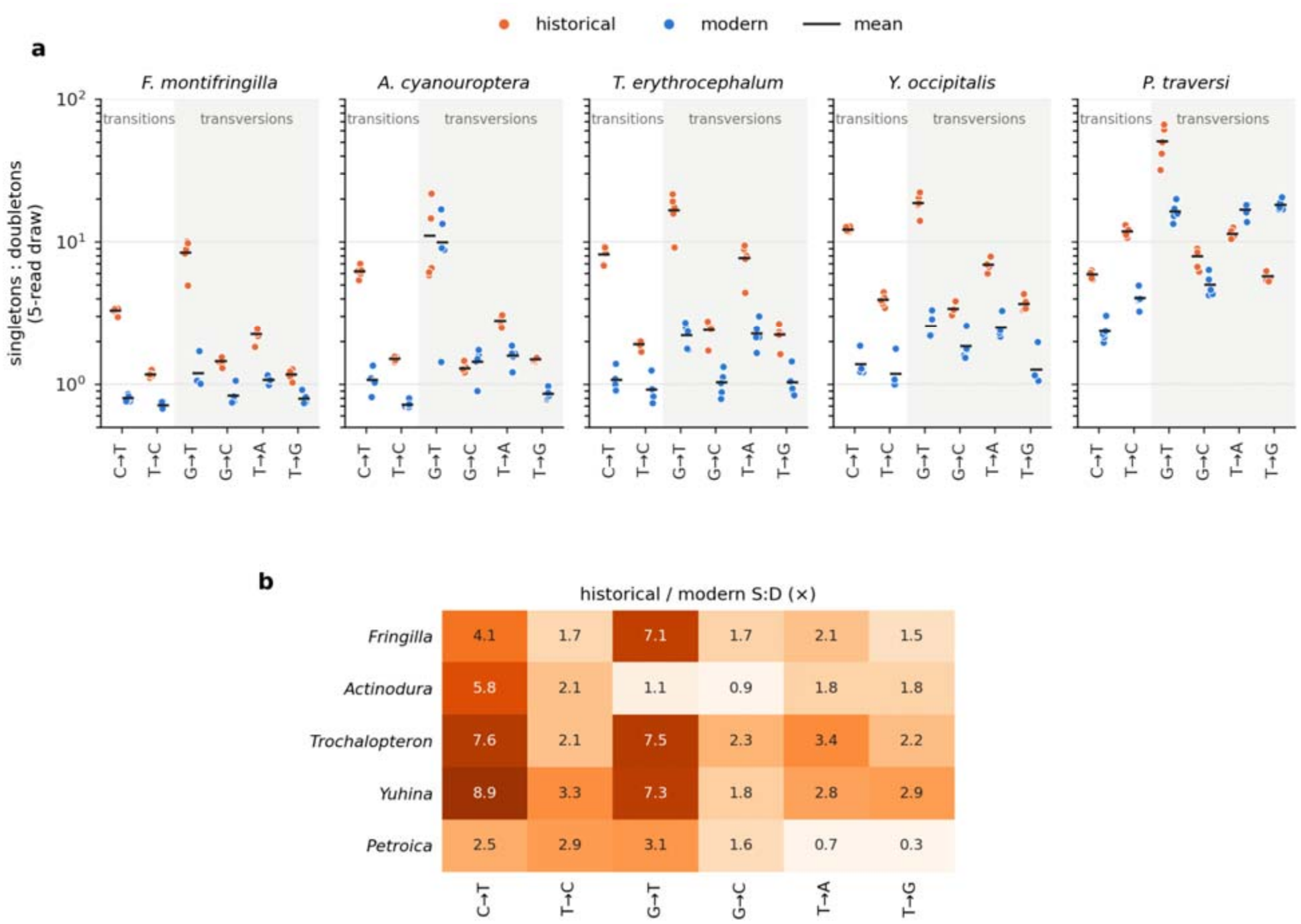
Excess single-read substitutions in historical samples. Each site was downsampled to five reads (the HEARTY draw), and non-reference bases were counted as singletons (one read) or doubletons (two identical reads); complementary substitutions are pooled. (a) Singleton:doubleton ratio per substitution class, per individual (bars: means). (b) Historical/modern fold change of the ratio (species means). Historical samples show an excess of single-read G→T transversions of similar size to the C→T deamination signal. The modern *A. cyanouroptera* libraries CRR1356156–9 show a comparable G→T excess.

